# The *C. elegans* ASPP homolog APE-1 is a junctional protein phosphatase 1 modulator

**DOI:** 10.1101/2022.02.15.480428

**Authors:** Gwendolyn M. Beacham, Derek T. Wei, Erika Beyrent, Ying Zhang, Jian Zheng, Mari M. K. Camacho, Laurence Florens, Gunther Hollopeter

## Abstract

How serine/threonine phosphatases are spatially and temporally tuned by regulatory subunits is a fundamental question in cell biology. Ankyrin repeat, SH3 domain, proline-rich-region-containing proteins (ASPPs) are protein phosphatase 1 (PP1) binding partners associated with cardiocutaneous diseases. ASPPs localize PP1 to cell-cell junctions, but how ASPPs localize and whether they regulate PP1 activity *in vivo* is unclear. Through a *C. elegans* genetic screen, we find that loss of the ASPP homolog, APE-1, suppresses a pathology called ‘jowls,’ providing us with an *in vivo* assay for APE-1 activity. Using structure-function analysis, we discover that APE-1’s N-terminal half directs the APE-1–PP1 complex to intercellular junctions. Additionally, we isolated mutations in highly conserved residues of APE-1’s ankyrin repeats that suppress jowls yet do not preclude PP1 binding, implying ASPPs do more than simply localize PP1. Indeed, *in vivo* reconstitution of APE-1 suggests the ankyrin repeats modulate phosphatase output, a function we find to be conserved among vertebrate homologs.

## Introduction

Serine/threonine phosphorylation is a reversible post-translational modification that regulates countless proteins throughout many eukaryotic cellular pathways (Olsen *et al*., 2006; Sharma *et al*., 2014). Genomes contain hundreds of kinases that orchestrate specific phosphorylation of protein substrates (Manning *et al*., 2002) yet the opposing dephosphorylation reactions are facilitated by only tens of phosphatase catalytic subunits (Shi, 2009; Chen, Dixon and Manning, 2017). This imbalance poses a conundrum: How do phosphatases achieve their specificity?

The major serine/threonine protein phosphatases, <u>P</u>rotein <u>P</u>hosphatase-<u>1</u> (PP1) and -<u>2A</u> (PP2A), are thought to acquire specificity by associating with non-catalytic regulatory subunits. These binding partners tune phosphatase activity through (1) *localization* of the catalytic subunit to a particular subcellular location, thereby concentrating phosphatase activity in a specific place and/or (2) *modulation* of the catalytic subunit to increase its activity toward some substrates and reduce its activity toward others (Hubbard and Cohen, 1993; Cohen, 2002; Bollen *et al*., 2010; Bertolotti, 2018). Classic examples include a glycogen-binding regulatory subunit that enhances PP1 catalytic subunit (hereafter referred to as PP1c) activity toward glycogen-metabolizing enzymes (Hubbard and Cohen, 1989), and a myosin-binding regulatory subunit that promotes PP1c dephosphorylation of myosins (Alessi *et al*., 1992). Through binding different regulatory subunits, a single phosphatase catalytic subunit can participate in potentially hundreds (Hendrickx *et al*., 2009) of distinct ‘holoenzymes.’ Yet for many of these holoenzymes, we still do not understand their functions, mechanisms, or *in vivo* targets (Bertolotti, 2018).

The <u>A</u>nkyrin repeat, <u>S</u>rc Homology 3 domain (SH3) domain <u>P</u>roline-rich-region-containing <u>P</u>roteins (ASPPs) are a conserved family of PP1c binding partners. ASPPs were originally characterized as p53 regulators (Iwabuchi *et al*., 1994; Samuels-Lev *et al*., 2001; Bergamaschi *et al*., 2003), but they have since been shown to bind PP1c with ∼100-fold greater affinity (Helps *et al*., 1995; Tidow *et al*., 2006; Patel *et al*., 2008; Llanos *et al*., 2011; Skene-Arnold *et al*., 2013). In vertebrates, the family includes ASPP1, ASPP2, and iASPP (Samuels-Lev *et al*., 2001; Sullivan and Lu, 2007). In mice, ASPP1 null mutants exhibit subcutaneous edema and mild lymphatic defects (Hirashima *et al*., 2008) while loss of ASPP2 results in postnatal lethality (Vives *et al*., 2006). Loss of iASPP, which is believed to be the most conserved family member (Bergamaschi *et al*., 2003), causes cardiocutaneous diseases in cows (Whittington and Cook, 1988; Simpson *et al*., 2009), mice (Herron *et al*., 2005; Toonen, Liang and Sidjanin, 2012), and humans (Falik-Zaccai *et al*., 2017). The molecular pathways underlying these pathologies remain unclear. One model proposes that disease states resulting from iASPP mutations stem from disruptions to desmosome junctions (Notari *et al*., 2015; Dedeić *et al*., 2018), yet how loss of iASPP perturbs these junctions is not well understood. Because ASPPs have been observed at intercellular junctions (Langton *et al*., 2009; Cong *et al*., 2010; Sottocornola *et al*., 2010; Dedeić *et al*., 2018), one proposed function for this protein family is to recruit phosphatase activity to these cellular locales (Royer *et al*., 2014; Notari *et al*., 2015; Bertran *et al*., 2019). While ASPPs appear to bind PP1c via their highly conserved C-terminal ankyrin repeats and SH3 domain, we do not yet understand how the ASPPs themselves localize to cellular junctions. Likewise, whether ASPPs simply recruit PP1c, or whether they also modulate phosphatase output remains unclear. *In vitro* phosphatase assays suggest that the ASPP C-terminal domains alter PP1c activity toward model substrates (Helps *et al*., 1995; Zhou *et al*., 2019) but whether these assays correspond to a function *in vivo* has remained challenging to explore.

Here we capitalize on a distinctive *C. elegans* pathology to delineate how ASPPs work *in vivo*. We call this phenotype ‘jowls,’ due to its hallmark presentation of bilateral, anterior bulges in the cuticle—the apical extracellular matrix secreted by epidermal cells that coats nematodes. Previously, jowls have been attributed to loss-of-function mutations in the clathrin adaptor complex, AP2 (Gu *et al*., 2013; Hollopeter *et al*., 2014). In this study, we discover dominant, gain-of-function forms of the Inversin homolog, MLT-4 (Lienkamp, Ganner and Walz, 2012; Lažetić and Fay, 2017), that cause jowls. MLT-4 localizes to apical junctions of epidermal cells where it is believed to regulate molting—the process by which larvae replace their old cuticle with a new one (Lažetić and Fay, 2017). MLT-4 has also been linked genetically to AP2-dependent trafficking in the epidermis (Joseph *et al*., 2020). We performed a genetic screen for suppressors of the gain-of-function forms of MLT-4, but instead of clarifying the connection between MLT-4 and AP2, this screen revealed a novel, and unexpected, regulator of jowls: APE-1, the sole *C. elegans* homolog of the ASPPs. Thus, while the biology underlying jowls remains unclear, we utilized this phenotype as an *in vivo* assay for APE-1 function.

Using unbiased proteomics, we find that like the vertebrate and *Drosophila* homologs, APE-1 binds a PP1c called GSP-2. Both APE-1 and GSP-2 localize to cell-cell junctions in the epidermis. Through structure-function analysis, we then discover that the N-terminal half of APE-1 dictates localization of the APE-1–GSP-2 complex to junctions. Additionally, we find that APE-1 does more than simply recruit GSP-2 to junctions: Characterization of the missense mutations isolated in our genetic screen coupled with reconstitution studies of APE-1 function *in vivo* reveals that the highly conserved ankyrin repeats modulate phosphatase output. We propose ASPPs have targeting modules for intercellular junctions encoded within their N-terminal region, and specify dephosphorylation reactions at these cellular sites via their C-terminal domains.

## Results and Discussion

### Dominant mutations in *mlt-4* cause jowls

In a mutagenesis screen for jowls (Hollopeter *et al*., 2014), we isolated a dominant, gain-of-function mutation (E470K) in the Inversin homolog MLT-4. We introduced this point mutation *de novo* using CRISPR and verified that it caused jowls (Figure 1A, second from left). These mutant worms will be referred to as *mlt-4(E470K)*. Separately, we tagged wildtype MLT-4 at its endogenous C-terminus with a bipartite TagRFP:HA-tag (MLT-4:RFP) and found that this modification also caused jowls (Figure 1A, center left). We will refer to these animals as *mlt-4:RFP*. Curiously, the effect of the RFP tag was more severe: *mlt-4(E470K)* are slightly ‘dumpy’ (shorter than wildtype worms), while *mlt-4:RFP* are even dumpier (Supplemental Figure 1A & B).

**Figure 1.**
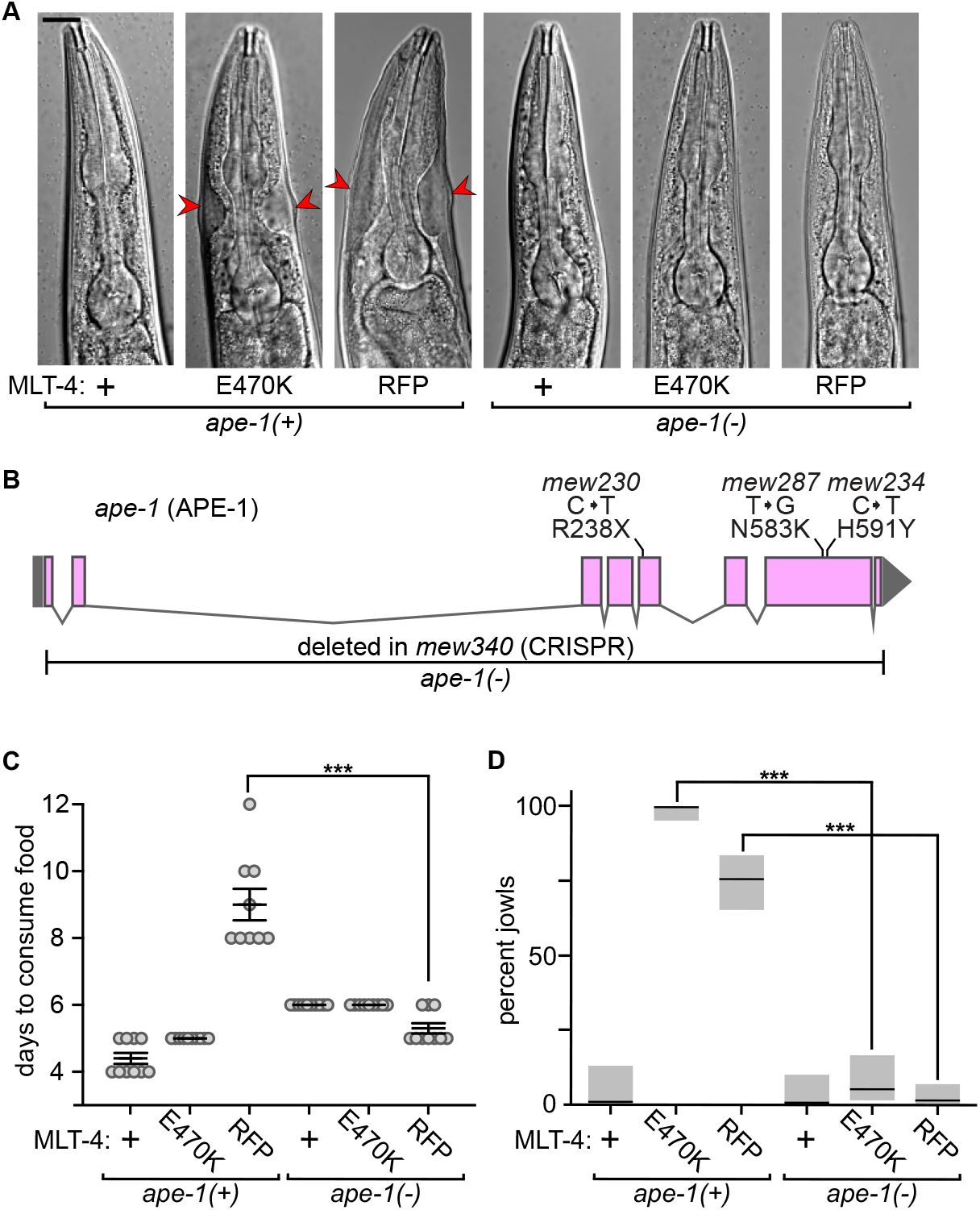
Loss of *ape-1* suppresses dominant *mlt-4* jowls alleles. **A**. Representative images of worm heads. Arrowheads indicate jowls; anterior is up; scale bar is 20 μm. **B**. Model of the *ape-1* locus. Boxes represent exons. Mutations isolated in the *mlt-4:RFP* suppressor screen are indicated above. Deletion allele used throughout this study, *ape-1(-)*, is indicated below. **C**. Fitness assay. Data points indicate days required for worms to multiply and exhaust food supply on culture plates, n = 9–10 plates. Bars represent mean ± SEM. **D**. Jowls assay. Bars indicate percentage of worms exhibiting jowls, n = 47–159 worms. Gray boxes represent 95% confidence intervals. ***p<0.0001, ANOVA analysis with Tukey’s post-hoc test. +, wildtype; -, knockout; E470K, single amino acid change; RFP, C-terminal TagRFP:HA-tag.

### Loss of *ape-1* suppresses jowls

To identify components downstream of MLT-4, we performed a suppressor screen. We mutagenized the more severe *mlt-4:RFP* with *N*-ethyl-*N*-nitrosourea (ENU) and over multiple generations, the animals that acquired mutations conferring increased fitness outcompeted sick *mlt-4:RFP* siblings on the culture plates. We singled suppressed hermaphrodites, one from each plate, for subsequent complementation analysis. Three of the isolated suppressors (one recessive and two semidominant) were in a single genetic complementation group. Using the sibling subtraction method (Joseph, Blouin and Fay, 2018) and whole genome sequencing, we found that all three of these strains contained mutations in the gene *ape-1* (protein APE-1). One was an early stop codon (R238X, recessive) and the other two were missense mutations (N583K, H591Y, both semidominant) (Figure 1B).

To confirm that jowls are APE-1-dependent, we generated a homozygous deletion of *ape-1* using CRISPR (Figure 1B) and crossed this null allele into both *mlt-4:RFP* and *mlt-4(E470K)*. Deletion of *ape-1* indeed suppressed the fitness defect of *mlt-4:RFP* as measured by the number of days required for a population of worms to expand and consume their bacterial food source on culture plates (Figure 1C). Note that in this assay, *ape-1* knockout animals exhibited a slight fitness defect and we did not detect suppression of *mlt-4(E470K)* (Figure 1C). However, when we scored the number of adult worms exhibiting jowls from a synchronized population, it was clear that loss of *ape-1* suppressed the jowls caused by both gain-of-function forms of MLT-4 (Figure 1D; Figure 1A, representative images). Additionally, knocking out *ape-1* suppressed the dumpy phenotype of *mlt-4:RFP* (Supplemental Figure 1B).

We found that loss of *ape-1* also suppressed the jowls of previously characterized mutants lacking either the AP2 activator, FCHO-1, or one of the AP2 subunits, APA-2 (Gu *et al*., 2013; Hollopeter *et al*., 2014; Beacham *et al*., 2018) (Supplemental Figure 2). Therefore, APE-1 is required for the jowls phenotype in general, not just for jowls caused by the gain-of-function forms of MLT-4. Although we do not yet fully understand the biology underlying the jowls phenotype, we capitalized on the simplicity of this phenotype as an *in vivo* assay for APE-1 activity. Unless noted, all subsequent assays were performed in the *mlt-4:RFP* genetic background.

### APE-1 forms a complex with GSP-2

To identify APE-1’s binding partners, we immunoprecipitated APE-1 endogenously tagged with a 3xFLAG:GFP (APE-1:GFP) from whole-worm lysates and analyzed the entire elution with mass spectrometry using <u>Mu</u>lti<u>d</u>imensional <u>P</u>rotein Identification <u>T</u>echnology (MudPIT) (Washburn, Wolters and Yates, 2001). The top interacting partner was the PP1c called GSP-2 (Figure 2A; Supplemental Table 1). Notably, p53, another characterized binding partner of ASPPs (Iwabuchi *et al*., 1994), was not detected. Nor was MLT-4, implying that APE-1 and MLT-4 exist in separate complexes (Supplemental Table 1). We performed a similar experiment in transiently transfected tissue culture cells (HEK293T) using HaloTagged vertebrate ASPPs (the APE-1 homologs) as baits. Consistent with our results from the APE-1 immunoprecipitations, the top hits consisted of PP1c isoforms and none of the ASPPs precipitated p53 (Figure 2B; Supplemental Table 2).

**Figure 2.**
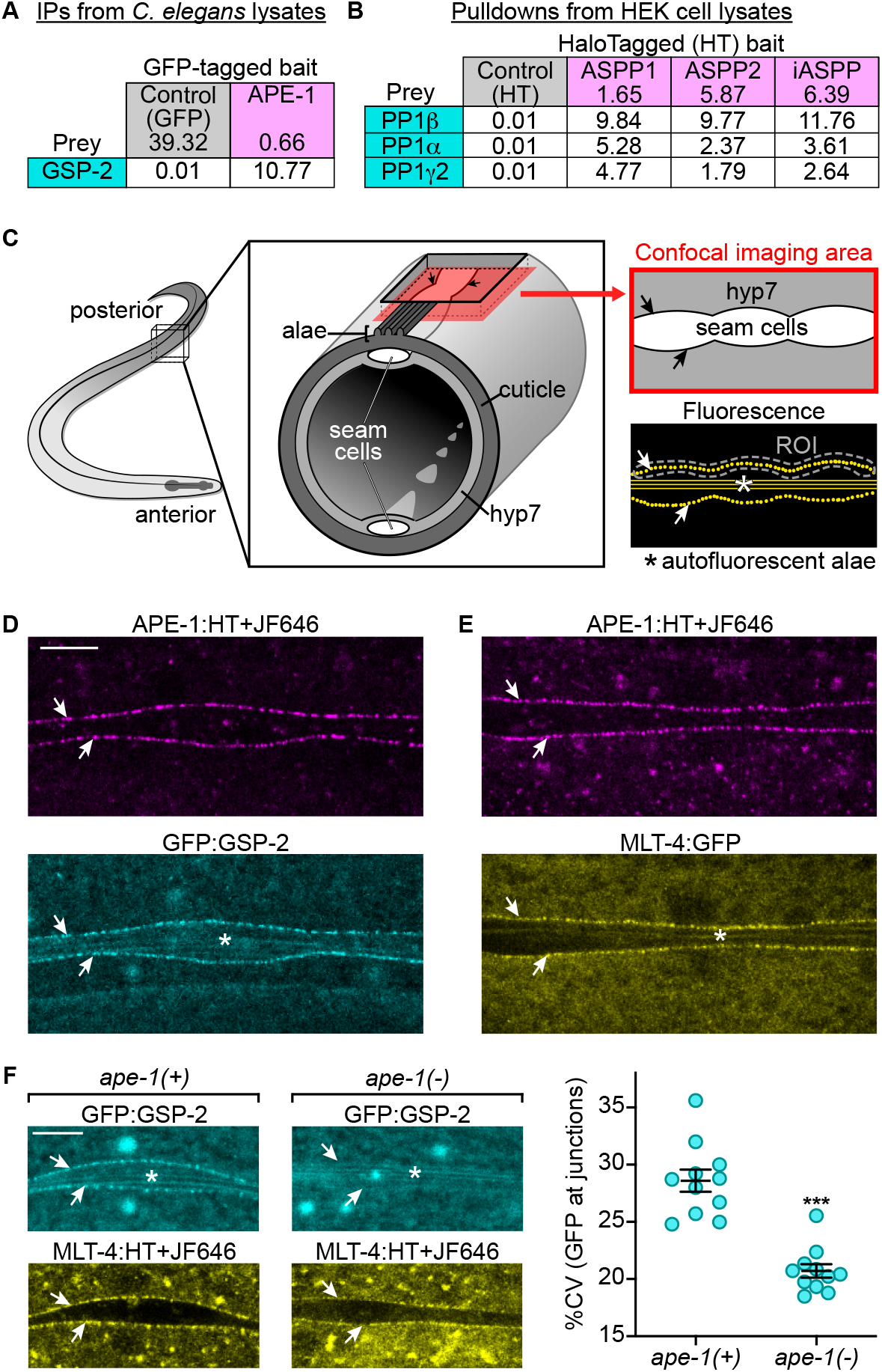
APE-1/ASPPs form a complex with GSP-2/PP1c. **A & B**. <u>Mu</u>lti<u>d</u>imensional <u>P</u>rotein Identification <u>T</u>echnology (MudPIT) mass spectrometry analysis of APE-1:GFP:3xFLAG immunoprecipitations (IPs) from whole *C. elegans* lysates (A) and of ASPP:HT pulldowns from HEK293T cell lysates (B). Values are the relative abundance of each protein in the sample as estimated by <u>d</u>istributed <u>N</u>ormalized <u>S</u>pectral <u>A</u>bundance <u>F</u>actor (dNSAF x 100, %dNSAF). **C**. Diagram depicting the adult worm epidermis and cuticle. The region imaged in D-F is indicated (red square) with the corresponding fluorescent image (right, bottom). Arrows point to junctions. White asterisks mark autofluorescent alae. Example region of interest (ROI, gray dashed line) used for measuring pixel intensity variation in subsequent experiments is drawn. **D-F**. Confocal images of endogenously-tagged APE-1 (magenta), GSP-2 (cyan), and MLT-4 (yellow) in the epidermis of adult worms. HaloTag was labeled with a chemical dye (HT+JF646) prior to imaging. **F**. Worms were *ape-1(+)* or *ape-1(-)* (indicated above). Left: Note nuclear localized GFP:GSP-2. Right: Variation in GFP:GSP-2 pixel intensities (percent coefficient of variance, %CV) in a region of interest (ROI, schematic Figure 2C) circumscribing junctional MLT-4:HT, n = 11 worms. Bars represent mean ± SEM. ***p<0.0001, compared to *ape-1(+)*, unpaired Student’s t-test. All scale bars are 10 μm.

Our results are consistent with previous reports that the highly conserved C-terminal ankyrin repeats and SH3 domain of ASPPs bind PP1c (Helps *et al*., 1995; Bertran *et al*., 2019; Zhou *et al*., 2019). Interestingly, APE-1 precipitated only one PP1c isoform (GSP-2) from worm lysates while the ASPPs precipitated three PP1c isoforms (PP1*α*, PP1*β*, and PP1*γ*2) from tissue culture cell lysates. All PP1c isoforms precipitated by ASPPs have a polyproline SH3 domain-binding motif (PxxPxR) near their C-termini that has been shown to be important for binding ASPPs. Within this polyproline motif is a cyclin-dependent kinase phosphorylation site that might further regulate binding of PP1c to ASPPs (Skene-Arnold *et al*., 2013). By contrast, GSP-2 is the only *C. elegans* PP1c with this C-terminal polyproline motif, which might explain why APE-1 did not precipitate the other widely expressed PP1c homolog, GSP-1 (Sassa *et al*., 2003; Wu *et al*., 2012). These results support the hypothesis that sequences in the C-termini of PP1c specify binding to ASPPs (Bertran *et al*., 2019).

### APE-1 and GSP-2 localize to epidermal junctions

To determine where APE-1 and GSP-2 co-localize *in vivo*, we endogenously tagged APE-1 on its C-terminus with a myc-tag:HaloTag (APE-1:HT), and GSP-2 on its N-terminus with a 3XFLAG:GFP (GFP:GSP-2), which has previously been shown to be functional (Hattersley *et al*., 2016). We then fed the worms a chemical dye (JF646) to covalently label the HaloTag (Los *et al*., 2008; Encell, 2013; Grimm *et al*., 2015) and imaged them using confocal microscopy.

While APE-1 and GSP-2 were observed in multiple different tissues, they were both present in the epidermis where the jowls phenotype originates (Gu *et al*., 2013). Here they localized to repetitive punctate structures lining the cell-cell junctions between the two major epidermal syncytia: hyp7 and the lateral seam cells (schematic Figure 2C; Figure 2D) (Chisholm and Hsiao, 2012). GSP-2 also localized to nuclei, consistent with previous studies (Hattersley *et al*., 2016). Interestingly, MLT-4 localizes to puncta at apical epidermal junctions (Lažetić and Fay, 2017) that appear to be similar to the junctional puncta to which APE-1 and GSP-2 localize (Figure 2E). This subcellular localization of APE-1, GSP-2, and MLT-4 might underlie the genesis of the jowls phenotype. Notably, the AP2 clathrin adaptor subunit, APA-2, has also been reported at these junctions in larval animals (Hadwiger *et al*., 2010). However, it is unclear whether AP2 or MLT-4 act in the same pathway as APE-1 and GSP-2.

PP1c has been observed at tight junctions in cultured canine kidney cells (Traweger *et al*., 2008), and at apical junctions in *Drosophila* retinas where localization appears to be dependent upon ASPP and another binding partner, CCDC85 (Bertran *et al*., 2019). We wanted to determine whether in *C. elegans*, GSP-2 localization depends on APE-1. To mark the epidermal junctions independently of APE-1, we endogenously tagged MLT-4 on its C-terminus with a HaloTag (MLT-4:HT) and labeled it with JF646 prior to imaging. We used confocal microscopy to quantify the variation in pixel intensity of GSP-2 at the junctions as delimited by MLT-4. Note that MLT-4 junctional signal appears reduced in *ape-1(-)*. Junctional GSP-2 puncta were not apparent in the *ape-1* knockout, though GSP-2 nuclear localization (Figure 2F) and expression (Supplemental Figure 3) appeared unchanged. Therefore, localization of GSP-2 to epidermal junctions depends on APE-1.

### The APE-1 N-terminus localizes to junctions

How do ASPPs localize to cell-cell junctions? While the C-terminal domains of ASPP proteins are highly conserved, the N-terminal half of the proteins is less obviously so and a function for this region has been difficult to pinpoint. Curiously, overexpression of the ASPP2 N-terminus disrupts localization of endogenous ASPP2 in cultured cells (Cong *et al*., 2010), suggesting that this region of ASPPs might confer binding to intercellular junctions.

To determine which portions of APE-1 are necessary and sufficient to localize to the epidermal junctions, we edited the *ape-1* locus with CRISPR to generate GFP-tagged APE-1 fragments of either the C-terminal ankyrin repeats and SH3 domain (amino acids 520-769, hereafter called the C-terminus), or the N-terminal half (amino acids 1-519, hereafter called the N-terminus) (Figure 3A, top schematic). While the APE-1 C-terminus no longer localized to the epidermal junctions, the APE-1 N-terminus localized just as well as the full-length protein (Figure 3A, middle and bottom).

**Figure 3.**
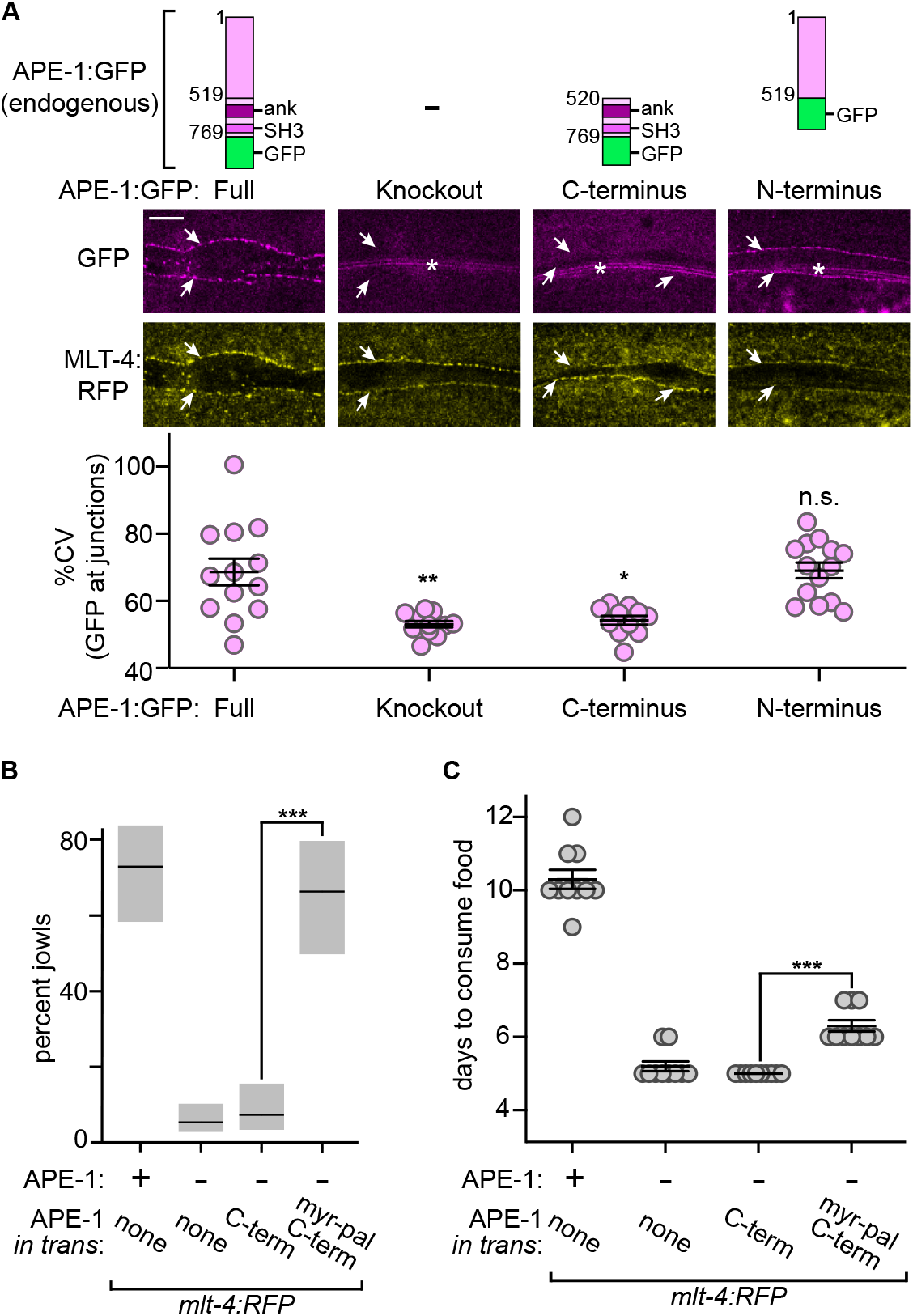
The N-terminus of APE-1 recruits the phosphatase complex to junctions. **A**. Top: Schematic of APE-1:GFP truncations. Numbers indicate amino acids. ank, ankyrin repeats; SH3, Src Homology 3 domain. Middle: Confocal slices in the epidermis of adult worms expressing APE-1:GFP truncations and MLT-4:RFP. Arrows point to junctions. White asterisks mark autofluorescent alae (see schematic Figure 2C). Scale bar is 10 μm. Bottom: Variation in GFP pixel intensities (percent coefficient of variance, %CV) in a region of interest (ROI, see schematic Figure 2C) circumscribing junctional MLT-4:RFP, n = 11–14 worms. **B**. Jowls assay. Bars indicate percentage of worms exhibiting jowls, n = 36–147 worms. Gray boxes represent 95% confidence intervals. **C**. Fitness assay. Data points indicate days required for worms to multiply and exhaust food supply on culture plates, n = 10 plates. Bars represent mean ± SEM. ***p<0.0001, **p<0.001, *p<0.01, n.s., not significant; compared to full APE-1:GFP (A) or as indicated (B and C), ANOVA analysis with Tukey’s post-hoc test. APE-1 C-termini were expressed *in trans* as single copy transgenes. C-term, C-terminus; myr-pal, myristoylation-palmitoylation signal; +, wildtype; -, knockout.

We tested if localizing the C-terminal phosphatase-binding domains to the membrane was sufficient to bypass the requirement for the N-terminus. To do this, we replaced APE-1’s N-terminus with a previously characterized myristoylation-palmitoylation (myr-pal) signal (Ramulu and Nathans, 2001). Indeed, membrane recruitment of the C-terminus restored jowls (Figure 3B), yet only partially restored the fitness defect of *mlt-4:RFP* (Figure 3C). In contrast, expression of the soluble C-terminus failed to restore either phenotype (Figure 3B & C). These data suggest the jowls phenotype is exquisitely sensitive to the output of the APE-1–GSP-2 complex whereas the fitness assay may be less so.

Because the membrane-localized C-terminus does not fully restore the fitness defect of *mlt-4:RFP* (Figure 3C), the N-terminal region of APE-1 likely localizes to a more specific cellular location. This observation is consistent with previous reports that the N-terminus of iASPP binds a desmosome protein, desmoplakin (Notari *et al*., 2015), while the N-terminus of ASPP2 binds a cell polarity regulator, PAR-3 (Cong *et al*., 2010; Sottocornola *et al*., 2010), which exhibits a junctional-localization pattern reminiscent of APE-1 in *C. elegans* (Castiglioni et al. 2020).

However, PAR-3 does not appear to be a direct binding partner of APE-1 according to our MudPIT analysis (Supplemental Table 1). Nonetheless, we have demonstrated that the N-terminus of APE-1 is sufficient to localize to intercellular junctions, revealing a clear activity for this part of the protein.

### Truncation of GSP-2 precludes interaction with APE-1 but does not suppress jowls

Structures of PP1c in complex with the C-terminal ankyrin repeats and SH3 domain from two different ASPPs have been determined (Bertran *et al*., 2019; Zhou *et al*., 2019). These structures reveal that the SH3 domain interacts with a PxxPxR motif in the C-terminus of PP1c. Immediately following this motif in PP1c are several conserved lysines which have been found to be important for binding the ASPPs (Bertran *et al*., 2019). To probe the APE-1–GSP-2 interaction, we truncated the C-terminal 13 amino acids of GSP-2, which removed these key lysines (Figure 4A, Δ13 indicated with a bracket). Indeed, when we immunoprecipitated GSP-2(Δ13) from whole-worm lysates, we no longer detected APE-1 in the elution sample (Figure 4B). We then used confocal microscopy to query localization of GSP-2(Δ13). Consistent with our biochemical experiments, GSP-2(Δ13) was absent from the epidermal junctions (Figure 4C) despite being stably expressed (Supplemental Figure 3).

**Figure 4.**
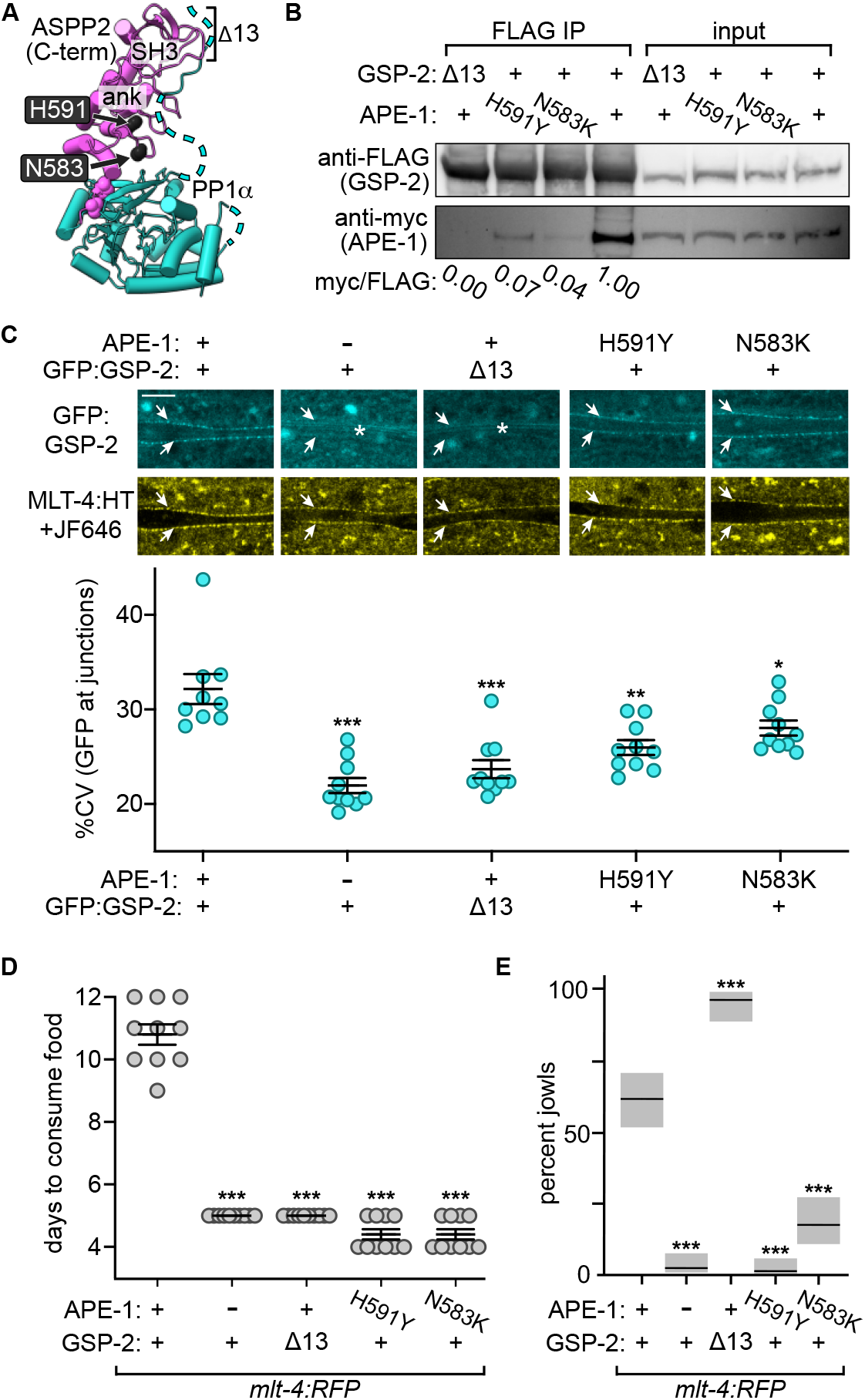
Perturbations of the APE-1–GSP-2 complex disrupt GSP-2 localization and suppress *mlt-4:RFP*. **A**. Predicted location of the mutated APE-1 residues (worm numbers N583 and H591) in the crystal structure of *H. sapiens* ASPP2 C-terminal domains (C-term) in complex with PP1*α* (PDB ID: 6GHM). ASPP2 is magenta, PP1*α* is cyan. Dashed lines represent unstructured regions. Residues deleted in GSP-2(Δ13) are indicated (bracket, Δ13). ank, ankyrin repeats; SH3, Src Homology 3 domain. **B**. Western blot analysis to detect a myc epitope on APE-1:HT proteins that co-precipitated with 3xFLAG:GFP:GSP-2 from worm lysates. Anti-myc to anti-FLAG intensity ratios were determined and normalized to the APE-1(+) GSP-2(+) ratio (values below). **C**. Confocal slices in the epidermis of adult worms expressing GFP:GSP-2 and MLT-4:HT, stained with JF646. Arrows point to junctions. White asterisks mark autofluorescent alae (see schematic Figure 2C). Scale bar is 10 μm. Bottom: Variation in GFP:GSP-2 pixel intensities (percent coefficient of variance, %CV) in a region of interest (ROI, schematic Figure 2C) circumscribing junctional MLT-4:HT, n = 9–10 worms. Bars represent mean ± SEM. **D**. Fitness assay. Data points indicate days required for worms to multiply and exhaust food supply on culture plates, n = 10 plates. Bars represent mean ± SEM. **E**. Jowls assay. Bars indicate percentage of worms exhibiting jowls, n = 78–127. Gray boxes represent 95% confidence intervals. *p<0.05, **p<0.001; ***p<0.0001, compared to strains with wildtype (+) versions of APE-1 and GSP-2, ANOVA analysis with Tukey’s post-hoc test. +, wildtype version; - knockout; HT, HaloTag; N583K and H591Y, single amino acid changes; Δ13, truncation of the final 13 amino acids.

To determine if this interface is required for a functional APE-1–GSP-2 complex *in vivo*, we generated GSP-2(Δ13) in *mlt-4:RFP*. Despite having increased fitness (Figure 4D) consistent with disruption of the APE-1–GSP-2 complex (Figure 4B & C), these animals still had jowls (Figure 4E) implying that APE-1 might transiently act upon stray GSP-2(Δ13) molecules that encounter the junctions. This residual activity could be mediated by the remaining PxxPxR motif in GSP-2, the RVxF motif in APE-1 that has been shown to bind PP1c in other systems (Skene-Arnold *et al*., 2013; Bertran *et al*., 2019), or perhaps APE-1’s poorly understood ankyrin repeats (Zhou *et al*., 2019).

### APE-1 missense mutations suppress jowls yet retain residual GSP-2 binding

Because the missense mutations we isolated in the *mlt-4:RFP* suppressor screen are in the highly conserved ASPP ankyrin repeats (Supplemental Figure 4A), we mapped them onto the crystal structures and found they are near the interface with PP1c (Figure 4A). To test if the missense mutations perturb the complex, we transiently transfected HEK293T cells with HaloTagged *H. sapiens* iASPP C-terminal domains in which the homologous residues were mutated (N657K and H665Y). Indeed, these mutant proteins pulled-down ∼50% less PP1c, as detected by western blot (Supplemental Figure 4B). When we compared the amount of APE-1 bound to GSP-2 in worm lysates, we detected less of the mutant APE-1 proteins compared to the wildtype, but more than was detected for GSP-2(Δ13) (Figure 4B). By microscopy, GSP-2 was still localized to the epidermal junctions in these mutants, yet the puncta were slightly less bright (Figure 4C). In contrast to the partial suppression observed for GSP-2(Δ13), the missense mutations suppressed both the fitness defect (Figure 4D) and jowls (Figure 4E) of *mlt-4:RFP*. Thus, even though these mutant APE-1 proteins still bind residual GSP-2 (Figure 4B & C), they appear to be inactive. Therefore, in addition to mediating binding to GSP-2, APE-1’s ankyrin repeats might confer an activity that exerts an additional layer of control over the phosphatase.

### The APE-1 C-terminus modulates GSP-2

To determine whether the ankyrin repeats might indeed influence phosphatase activity, we tested whether localizing GSP-2 in the absence of the APE-1 C-terminus (the ankyrin repeats and SH3 domain) was sufficient to cause jowls in the *mlt-4:RFP* background. Because the APE-1 N-terminus is sufficient to localize to junctions (Figure 3A), we directly fused GSP-2 to this sequence, thereby simultaneously removing the APE-1 C-terminus (Figure 5A, top schematic, third from left). We reasoned the following: If the only function of the APE-1 C-terminus is to bind GSP-2, this APE-1 N-terminus:GSP-2 fusion would be sufficient to cause jowls. By contrast, if the APE-1 C-terminus is also required to modulate GSP-2, this fusion protein would be insufficient for jowls. Consistent with the model that the APE-1 C-terminus might confer additional activity, APE-1 N-terminus:GSP-2 failed to restore jowls (Figure 5A, 3rd from left) unless the APE-1 C-terminus was also expressed *in trans* (Figure 5A, 4th from left). This recapitulation of APE-1 activity was blocked by either missense mutation in the ankyrin repeats (Figure 5A, 5th & 6th from left) or, if GSP-2 was removed from the APE-1 N-terminus (Figure 5A, 7th from left) implying that jowls require both GSP-2 localization to epidermal junctions and additional regulation by the APE-1 C-terminus. This activity conferred by the C-terminus appears to be conserved as expression of the homologous fragment from either mouse ASPP1 (amino acids 847-1087) or ASPP2 (amino acids 902-1134) was also sufficient to cause jowls (Figure 5B).

**Figure 5.**
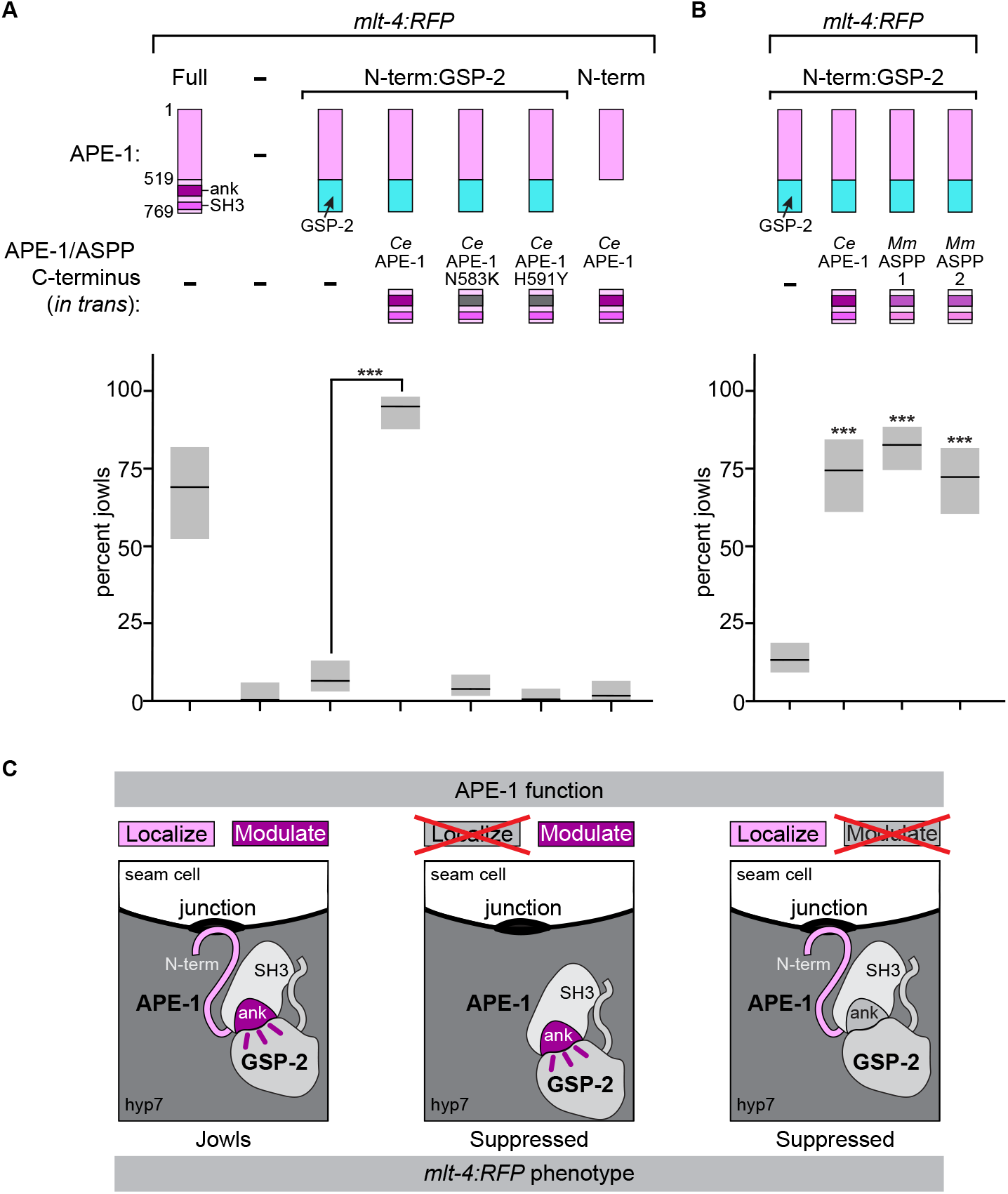
C-terminal domains from APE-1 or ASPP modulate GSP-2. **A & B**. Jowls assay. Bars indicate percentage of worms exhibiting jowls, n = 36–191. Gray boxes represent 95% confidence intervals. ***p<0.0001, ANOVA analysis with Tukey’s post-hoc test, in B compared to no C-terminus (-). Note that proteins expressed from the *ape-1* locus (top) had a GFP tag (not represented in schematic) and APE-1/ASPP C-terminal domains (middle) were expressed *in trans* as single copy transgenes. **C**. Model of the junctional phosphatase complex highlighting the new activities of APE-1 we identified. The N-terminus dictates localization, while the ankyrin repeats regulate the phosphatase. The *mlt-4:RFP* phenotype is suppressed when these functions are perturbed. -, absent; ank, ankyrin; SH3, Src Homology 3 domain; N-term, N-terminus; *Ce, C. elegans*; *Mm, M. musculus*.

Classically, phosphatase regulatory subunits are thought to increase phosphatase activity toward desired substrates and reduce activity toward other targets (Hubbard and Cohen, 1993; Bertolotti, 2018). Numerous targets have been proposed for ASPP–PP1 holoenzymes, including transcription cofactors in the Hippo pathway (Liu *et al*., 2011; Royer *et al*., 2014), p53 (Zhou *et al*., 2019), and desmoplakin (Notari *et al*., 2015). However, it is not clear if any of these underlie the jowls phenotype. We can speculate about potential targets based on APE-1 binding partners identified in our MudPIT analysis. Several of the top hits were myosin light and heavy chains (Supplemental Table 1). Intriguingly, iASPP was recently demonstrated to bind the myosin Myo1c (Mangon *et al*., 2021) and the ASPP-related myosin phosphatase regulatory subunit MYPT1 binds PP1c via ankyrin repeats (Tanaka *et al*., 1998; Terrak *et al*., 2004). Other top hits were tubulin and actin (Supplemental Table 1), consistent with previous reports that iASPP associates with the tubulin-binding protein EB1 (Mangon *et al*., 2021). Notably, the hyp7-seam cell junctions are enriched for cytoskeletal proteins during molting (Costa, Draper and Priess, 1997; Lažetić *et al*., 2018). While there are several known phosphorylation sites on both tubulin and actin (Wloga, Joachimiak and Fabczak, 2017; Varland, Vandekerckhove and Drazic, 2019), it is not clear if any are the target of the APE-1–GSP-2 holoenzyme.

In conclusion, we have demonstrated that the *C. elegans* ASPP homolog, APE-1, is required for jowls—a phenotype caused by dominant mutations in the molting regulator, MLT-4, or recessive mutations in the clathrin adaptor complex, AP2. Pairing this *in vivo* readout for APE-1 activity with structure-function analysis enabled us to define two of APE-1’s activities as a phosphatase regulator: (1) APE-1’s localization to intercellular junctions is dictated by its N-terminal half and (2) APE-1’s highly conserved ankyrin repeats confer control over phosphatase output (Figure 5C). Identification of the relevant APE-1–GSP-2 phosphatase substrate will be required to delineate how the phosphatase is modulated.

## Materials and Methods

### Strains

*C. elegans* were maintained at room temperature (22–23 °C) on nematode growth medium (NGM) plates seeded with OP50 *E. coli*. Supplemental File 1 lists all strains (A) and alleles (B).

### *mlt-4:RFP* suppressor screen

Late L4–young adult *mlt-4:RFP* mutants (GUN405) were mutagenized for 4 hr at 22 °C in 0.5 mM N-nitroso-N-ethylurea (ENU, Sigma Aldrich N3385). Worms were washed in M9 buffer (22 mM KH_2_HPO_4_, 42.3 mM Na_2_HPO_4_, 85.6 mM NaCl, 1 mM MgSO_4_) and distributed across NGM growth plates seeded with concentrated OP50 bacterial culture. Once the population of worms had expanded and consumed all of the OP50, an approximately 2 × 2 cm chunk from each plate was transferred to a fresh plate. This process was repeated 2 times to select for worms with increased fitness. One suppressed animal per culture plate was singled to establish a suppressed strain. Males were generated from suppressed strains (1 hr heatshock at 34 °C in a circulating water bath) and crossed to hermaphrodites of other suppressed strains to test for genetic complementation.

The Sibling Subtraction Method (Joseph, Blouin and Fay, 2018) was used for whole genome sequencing sample preparation and analysis. Males generated from 3 independent suppressed strains belonging to a single complementation group (GUN514, GUN437, and GUN441) were crossed to the un-mutagenized parental strain (GUN405). For each strain, 8 suppressed F2s and 8 non-suppressed F2s were singled and their homozygosity was confirmed by phenotype of the F3s. When these plates were confluent, worms were washed off the plates with M9 buffer, pelleted by centrifugation at 500 x g, and washed two times with M9 buffer. Worms were frozen in liquid nitrogen and stored at -80 °C prior to genomic DNA preparation.

Genomic DNA was prepared from the pooled suppressed worms and pooled non-suppressed worms for each strain. Worm pellets were lysed in 10 volumes of lysis buffer (0.1 M Tris-Cl pH 8.5, 0.1 M NaCl, 50 mM EDTA pH 8.0, 1% SDS) containing 400 μg/mL Proteinase K (New England Biolabs) and incubated at 65 °C for 1 hr with rotation at 300 RPM. An equal volume of phenol:chloroform:isoamyl (25:24:1, Saturated with 10mM Tris, pH 8.0, 1mM EDTA) (Millipore Sigma) was added to each sample and shaken vigorously to mix. The phase layers were separated by spinning the samples in a room temperature tabletop centrifuge for 5 min at 17000 x g. The remaining phenol was removed by extraction with chloroform:isoamyl (24:1). DNA was precipitated from the aqueous layer by addition of 1/10 volume of 3 M NaoAC and 2.2 vol of ice cold 100% ethanol, and spun at max speed at 4 °C. The DNA pellet was washed in 70% ice cold ethanol. To remove contaminating RNA, the pellet was dissolved in 200 μL 10 mM Tris, 1 mM EDTA (pH 8) (TE) containing 10 mg/mL RNase A (Thermo Scientific), and incubated at 37 °C for 1 hr. Genomic DNA was precipitated again, but spooled off the top instead of pelleted. DNA was washed 2x with 70% ethanol, pelleted, and resuspended in TE.

Whole genome sequencing was performed by Novogene. Paired-end 150 bp libraries were sequenced on an Illumina Novaseq 6000 at 50x coverage and reads were mapped to the WBcel235 *C. elegans* reference genome. The allele frequency (AF) of detected variants was defined as the # alternate reads/the # total reads. Variant filtering FA4 as defined previously (Joseph, Blouin and Fay, 2018) was applied so that a variant allele in the suppressed sample was defined as homozygous and unique if the AF in the suppressed sample was ≥0.9, and the AF for the non-suppressed sample was ≤0.1. Mapped reads in the regions containing these candidate unique, homozygous variants were visualized on the integrative genome browser (IGV, Broad Institute, http://software.broadinstitute.org/software/igv/home) to confirm that read depth was high and variants were indeed unique to the suppressed sample.

### Preparation of worms for microscopy

For brightfield microscopy, worms were mounted on 2% agarose pads in 3 μL of PBS pH 7.4 with 10 mM sodium azide (Figure 1A) or in 3 μL of M9 with 20 mM sodium azide (body length assay, Supplemental Figure 1). Images were taken within 15–30 min of slide preparation. For confocal fluorescence microscopy, worms were mounted on 8–10% agarose pads in 3 μL of a 0.5 μM polystyrene bead slurry (Polysciences, Warrington, PA) in PBS pH 7.4 (Kim *et al*., 2013).

### Fitness assay

The fitness assay was performed as previously described (‘starvation assay’) (Hollopeter *et al*., 2014) with slight modifications. L4 hermaphrodites were picked and cultured at room temperature (22–23 °C) overnight. The next day, these worms (now adults) were distributed across multiple (usually 10) plates per strain, 3 worms per plate. Plates were incubated at 22– 23 °C and checked daily until worms had consumed all of the food.

### Jowls assay

Adult hermaphrodite worms were distributed across two plates per strain and removed after laying eggs for 5 hrs. Once the offspring were adults, jowls were counted under a dissecting microscope. The number of worms counted per strain ranged from 36–191. The person scoring jowls was blinded to the strain identity. Plates were incubated at 22–23 °C throughout the assay.

### Body length assay

For each strain, gravid adults were selected and imaged using brightfield microscopy on a Keyence microscope with a 10x objective. Images were analyzed in Fiji (Schindelin *et al*., 2012). A user-defined line was drawn through the middle of the worm from anterior to posterior and length of the line was measured.

### Whole-worm immunoprecipitations

Worms for immunoprecipitations were cultured at room temperature (22–23 °C) on 15 cm NGM plates seeded with concentrated OP50 *E. coli* culture mixed with chicken egg.

Immunoprecipitations for MudPIT samples (Data in Figure 2A and Supplemental Table 1): When plates were confluent but not yet starved, the unsynchronized population of worms were harvested from culture plates using cold H150 buffer (50 mM HEPES pH 7.6 and 150 mM KCl). Worms were settled on ice and centrifuged at 1000 x g at 4 °C for 2 min. Supernatant was removed and worms were washed twice more with cold H150 buffer. Worms were lysed in 25 mM HEPES pH 7.6, 75 mM KCl, and 5% glycerol with protease inhibitors (Roche, 1 tablet per 10 mL lysis buffer). Worm slurry was frozen in liquid nitrogen and stored at -80 °C until lysis. Frozen worms were ground in a mortar and pestle chilled with liquid nitrogen. When no intact worms remained as verified under a dissecting scope, lysed worms were thawed on ice and diluted 5-fold with 50 mM HEPES pH 7.6, 150 mM KCl, and 10% glycerol containing 0.005% IGEPAL (CA-630, Sigma). Lysate was spun at 19650 x g for 20 min at 4 °C and the cleared supernatant was filtered in a 0.45 μm filter (Merck Millipore). M2 magnetic FLAG beads (Sigma-Aldrich, M8823) (100 μL of 50% slurry for every ∼2.5 g starting worm pellet) were equilibrated in lysis buffer and added to the filtered supernatant which was rotated with the beads overnight at 4 °C. Beads were pelleted with a magnet and the unbound supernatant was removed. Beads were washed with 20 packed bead volumes of lysis buffer twice, then once with TBS (Tris Ph 7.6 and 150 mM NaCl). Bound proteins were eluted with 5 packed bead volumes of 150 ng/μL 3xFLAG peptide (Sigma-Aldrich, F4799) in TBS for 30 min at 4 °C with rotation.

Immunoprecipitations in Figure 4B: These immunoprecipitations were performed as described above with the following modifications. Worms were harvested and washed in TBS, with centrifugations at 180 x g at 4 °C for 2 min. Worms were lysed in 20 mM Tris pH 7.6, 150 mM NaCl, 10% glycerol, 0.02% IGEPAL (CA-630, Sigma), 10 mM MgCl_2_, 1 mM CaCl_2_, 1X Protease Inhibitor Cocktail (Promega, #G6521), and 100 μg/mL DNase I (grade II from bovine pancreas, Roche). Frozen worms were lysed in a coffee grinder (Krups, F2034251) chilled with liquid nitrogen. The concentrated worm lysate was diluted 5-fold with 20 mM Tris pH 7.6, 150 mM NaCl, and 2% glycerol. Extra MgCl_2_, CaCl_2_, and DNase were added so their final concentrations were 22 mM MgCl_2_, 2.2 mM CaCl_2_, 120 μg/mL DNase. The lysate was spun and supernatant filtered as described above. Filtered lysates were incubated with the anti-FLAG beads (Sigma-Aldrich, M8823) for 2 hrs at room temperature (22 °C). Bound proteins were eluted with 125 ng/μL 3xFLAG peptide (Sigma-Aldrich, F4799) in TBS. 25% of these elution samples were separated by SDS-PAGE alongside 0.001% of input lysate samples for comparison.

### Tissue culture pulldowns

Mammalian vectors for expression of HaloTag fusion proteins were constructed using Gibson cloning (Gibson *et al*., 2009). A pcDNA5/frt vector encoding a 3’linker sequence with a TEV protease site and HaloTag was amplified with oDTW48-53 and recombined in a one-piece Gibson reaction to generate pDTW47. This construct was amplified with oDTW47-48 and recombined in 2-piece Gibson reactions with: (1) *M. musculus* ASPP1 (NM_011625.1) amplified from mouse brain cDNA with oDTW67-68 to generate pDTW50, (2) *M. musculus* ASPP2 (NM_173378.2) amplified from mouse brain cDNA with oDTW70-71 to generate pDTW51, and (3) a synthetic codon-optimized gBlock (IDT) encoding *H. sapiens* iASPP (NM_001142502.2) to generate pDTW53. pDTW53 was further amplified with oDTW261-262 and recombined in a one-piece Gibson reaction to generate pDTW81, which expressed amino acids 602-828 of *H. sapiens* iASPP with a C-terminal linker:TEV-site:HaloTag. One-piece Gibson reactions were used to insert single codon changes resulting in N657K and H665Y by amplifying pDTW81 with oDTW214-215 to generate pDTW94, and oDTW216-217 to generate pDTW87, respectively. Finally, a sequence encoding a short linker and HA-tag were fused to the 5’ end of the existing C-terminal linker, upstream of the TEV-site, in pDTW46, pDTW81, pDTW94, and pDTW87 by amplifying each with oDTW314-315 and recombining the amplicons in one-piece Gibson reactions. This generated pDTW117, pDTW123, pDTW125, and pDTW124, respectively. Supplemental File 1 lists all plasmids (C) and oligonucleotides (D). 150 mm dishes of HEK293T cells (∼80% confluent) were transfected with 10 μg of plasmid using 60 μg of linear Polyethylenimine 25 kDa (23966, Polysciences Inc) mixed in 2 mL Opti-MEM (31985070, ThermoFisher). 24 hr later, cells were scraped from dishes, collected by centrifugation and frozen at -80 °C. Pellets were lysed and bait proteins in the lysates were bound to HaloLink Magnetic Beads (Promega) according to manufacturer’s instructions. The beads were washed in TBS with 0.09% IGEPAL (CA-630, Sigma). Samples were incubated (2 hr at 21 °C with 1100 RPM shaking) with AcTEV protease (20 units in 100 μL, Invitrogen) to digest the protease site between the HaloTag and baits.

### MudPit analysis

Samples to be analyzed by MudPIT were precipitated with acetone as follows. The Tris concentration was adjusted to 100 mM, tris(2-carboxyethyl)phosphine (TCEP-HCl) (Pierce) was added to 10 mM, and samples were incubated at 55 °C for 1 hr. 2-chloroacetamide (Sigma-Aldrich) was added to 25 mM and samples were incubated for 30 min at room temperature, protected from light. Six volumes of pre-chilled acetone were added and precipitation was allowed to proceed overnight at -20 °C. The following day, samples were centrifuged at 8000 x g for 10 min at 4 °C. The acetone was decanted and the pellets were allowed to dry for a few minutes.

Protein samples were analyzed by MudPIT as described previously (Florens and Washburn, 2006). Briefly, precipitated proteins were resuspended in 30 uL of 100 mM Tris pH 8.5 with 8 M urea to denature proteins. Cysteines were reduced and alkylated prior to digestion with recombinant LysC and modified trypsin. Reactions were quenched by the addition of formic acid to the final concentration of 5%. After digestion, peptide samples were pressure-loaded onto 100 um fused silica microcapillary columns packed first with 9 cm of reverse phase material, followed by 3 cm of 5um Strong Cation Exchange material, followed by 1 cm of 5um C_18_ RP. The loaded microcapillary columns were placed in-line with a 1260 Quartenary HPLC. The application of a 2.5 kV distal voltage electrosprayed the eluting peptides directly into orbitrap Elite mass spectrometers equipped with a custom-made nano-LC electrospray ionization source. Full MS spectra were recorded on the eluting peptides over a 400 to 1600 *m*/*z* range, followed by fragmentation in the ion trap on the first to fifth most intense ions selected from the full MS spectrum. Dynamic exclusion was enabled for 90 s. Mass spectrometer scan functions and HPLC solvent gradients were controlled by the XCalibur data system.

RAW files were extracted into .ms2 file format using RawDistiller v. 1.0, in-house developed software. RawDistiller D(g, 6) settings were used to abstract MS1 scan profiles by Gaussian fitting and to implement dynamic offline lock mass using six background polydimethylcyclosiloxane ions as internal calibrants (Zhang *et al*., 2011). MS/MS spectra were first searched using ProLuCID with a mass tolerance of 10 ppm for peptide (Xu *et al*., 2015) and fragment ions. Trypsin specificity was imposed on both ends of candidate peptides during the search against the protein database. The human protein database contained 81592 human proteins (NCBI 2020-11-23 release), as well as 426 common contaminants such as human keratins, IgGs and proteolytic enzymes. The *C. elegans* protein database included 28,127 *C. elegans* proteins (NCBI 2020-05-30 release), as well as 426 common contaminants. To estimate false discovery rates (FDR), each protein sequence was randomized (keeping the same amino acid composition and length) and the resulting “shuffled” sequences were added to the database, for a total search space of 163,860 amino acid sequence for human datasets and 57,106 amino acid sequences for *C. elegans* datasets. Masses of 57.0215 Da were differentially added to cysteine residues to account for alkylation by CAM and 15.9949 Da were differentially added to methionine residues.

DTASelect v.1.9 was used to select and sort peptide/spectrum matches (PSMs) passing the following criteria set: PSMs were only retained if they had a DeltCn of at least 0.08; minimum XCorr values of 1.8 for singly-, 2.1 for doubly-, and 2.5 for triply-charged spectra; peptides had to be at least 7 amino acids long. Results from each sample were merged and compared using CONTRAST (Tabb, McDonald and Yates, 2002). Combining all replicate runs, proteins had to be detected by at least 2 peptides and/or 2 spectral counts. Proteins that were subsets of others were removed using the parsimony option in DTASelect on the proteins detected after merging all runs. Proteins that were identified by the same set of peptides (including at least one peptide unique to each protein group to distinguish between isoforms) were grouped together, and one accession number was arbitrarily considered as representative of each protein group. *NSAF7* was used to create the final reports on all detected peptides and non-redundant proteins identified across the different run (Zhang *et al*., 2010).The kite suite of software used to perform mass spectrometry data processing and analysis is archived in Zenodo (tzw-wen, 2022).

Raw data and search results files have been deposited to the Proteome Xchange (accession: PXD031435) via the MassIVE repository and may be accessed at ftp://MSV000088781@massive.ucsd.edu with password gunther-02-2022.

Mass Spectrometry data may also be accessed from the Stowers Original Data Repository (http://www.stowers.org/research/publications/libpb-1678).

### HaloTag ligand staining

Janelia Fluor 646 HaloTag Ligand (Promega) was added to 2.5 μM in OP50 bacterial culture on NGM plates. Mixed age worms were cultured on these plates overnight (13–19 hrs) at room temperature (22–23 °C) then washed using S buffer (6.45 mM K_2_HPO_4_, 43.55 mM KH_2_PO_4_, 100 mM NaCl) to seeded NGM plates lacking JF646 to ‘destain’ for 1–4 hr before imaging.

### Confocal microscopy

Adult worms were imaged on a Ziess LSM 880 confocal microscope (Biotechnology Resource Center, Cornell University, Ithaca, NY) with a 40x water immersion objective. Each set of strains was imaged in a single session using the same laser settings. Images were analyzed in Fiji (Schindelin *et al*., 2012).

For GFP:GSP-2 junctional pixel variance determination (Figure 2F; Figure 4C), images were acquired within 15 min of slide preparation. A confocal z-stack with its boundaries on either side of the MLT-4:HT+JF646 epidermal signal was acquired for each worm. The confocal slice with the most in-focus MLT-4:HT+646 signal at junctions was selected for quantification. A user-defined region of interest (ROI) circumscribing the MLT-4:HT+JF646 signal on one side of the hyp7-seam cell junctions was drawn. The mean intensity and standard deviation of GFP pixel intensities in the same confocal slice within the ROI were measured and used to calculate the coefficient of variance (standard deviation/mean). After drawing the ROI in the 646 channel, the user verified that neither GFP:GSP-2+ epidermal nuclei nor autofluorescent alae were present in the ROI in the GFP channel, since these were not the structures of interest to be quantified.

For the APE-1:GFP structure-function microscopy (Figure 3A), images were acquired within 15– 30 min of slide preparation. A confocal slice where MLT-4:RFP was most in-focus at the junctions was imaged. A user-defined ROI encircling the MLT-4:RFP signal along one side of the hyp7-seam cell junctions was drawn and the GFP coefficient of variance in the ROI was determined as described above.

### Western blots

Precast polyacrylamide gels (Bolt 4–12% Bis-Tris, Invitrogen) were used for all SDS-PAGE experiments. Samples were first denatured in 1X Bolt LDS Samples Buffer containing 12.5–25 mM dithiothreitol (DTT) and heated at 95 °C for 1 min. The Pierce Power Blot Cassette system (Thermo Scientific) was used to transfer proteins to PVDF Immobilon membranes (Merck Millipore). Blocking and antibody incubation steps were performed in EveryBlot Blocking Buffer (BioRad). Blots and gels were imaged using the Bio-Rad ChemiDoc MP system and band intensities were quantified using the associated ImageLab software.

Primary antibodies and dilutions included mouse anti-myc (1:1000, abcam, clone 9E10, ab32), mouse anti-FLAG (1:1000, sigma, clone M2, F3165), rabbit anti-beta actin (1:1000, abcam, ab8227), mouse anti-PP1 (1:1000, Santa Cruz, clone E-9, sc-7482), and rat anti-HA directly conjugated to HRP (1:500, clone 3F10, Roche). Secondary antibodies and dilutions included goat anti-mouse IRDye 800CW (1:20000, LI-COR, 926–32210) and goat anti-rabbit AlexaFlour 647 (1:2000, Life Technologies, A21245).

For Supplemental Figure 3, 20 adult worms were lysed in PBS pH 7.6 with 1X Bolt LDS Sample Buffer (Invitrogen) containing 25 mM dithiothreitol (DTT). Samples were frozen in liquid nitrogen and sonicated (1 s pulses at 70% amplitude for 3 min) in a cup horn (Branson Ultrasonics Corporation) chilled with circulating 4°C water. Samples were then heated for 10 min at 70 °C before SDS-PAGE.

### CRISPR-Cas9 transgenics

CRISPR-Cas9 edits were generated with ribonucleoprotein (RNP) complexes. The gonads of young adult hermaphrodites were injected with RNP mixes containing the following components: 0.25 μg/μL Cas9 (IDT 1081059), 0.1 μg/μL tracrRNA (IDT 1072534), 56 ng/μL gene-specific crRNA (IDT), 25 ng/μL gene-specific repair, and 40 ng/μL either the rol6(su1006) (pRF4) (Mello *et al*., 1991) or Prab-3::GFP:unc-54 3’UTR (pGH5) as a co-injection marker. Gene-specific repairs were generated and purified as described in (Ghanta and Mello, 2020), using unmodified oligos and Q5 high-fidelity polymerase (NEB) for the PCR reaction. After purification, repairs were incubated at 95 °C for 2 min, and then 10 seconds at 85 °C, 75 °C, 65 °C, 55 °C, 45 °C, 35 °C, 25 °C, and 12 °C. Repairs were kept on ice until they were added to the injection mix. F1 progeny from the 2 plates with the highest number of array-positive animals were moved to individual NGM plates and allowed to lay eggs for 2 days. Animals were then lysed in 10 mM Tris, 50 mM KCl, 1.5 mM MgCl_2_, 0.45% IGEPAL, 0.45% Tween-20, 0.8 U/μL Proteinase K (NEB), frozen at -80 °C, then incubated at 65 °C for 1 hr and 95 °C for 15 min. In order to identify correctly edited worms, PCR was used to amplify the target gene. For small inserts, a restriction site was engineered close to the edit. To confirm correct editing, the PCR product was sequenced.

### Data analysis

Data from fitness assays and microscopy were analyzed using one-way ANOVA tests with Tukey’s post-hoc analysis in GraphPad Prism (Version 9.2.0 for Mac, GraphPad Software, La Jolla, CA, USA, www.graphpad.com). For microscopy in Figure 2F, an unpaired, parametric two-tailed T-test was performed instead, also in GraphPad.

For the jowls assays, RStudio was used to fit a generalized linear model for a binomial distribution. In order to account for complete separation (which results when one or more categories consist of 100% jowls or 100% no jowls), logistic regression using Firth’s bias-correction was implemented using the brglm2 package. ANOVA analysis was performed using the emmeans package, adjusting for multiple comparisons using Tukey’s method.

### Molecular visualization

The structural representation in Figure 4A was prepared in ChimeraX (Goddard *et al*., 2018; Pettersen *et al*., 2021) (version 1.3 (2021-12-08) for Mac, https://www.rbvi.ucsf.edu/chimerax).

### Sequence alignment

The sequence alignment in Supplemental Figure 4A was performed using Jalview (Waterhouse *et al*., 2009).

## Additional Files

Supplementary File 1. A. Strains. B. Alleles. C. Plasmids. D. Oligonucleotides.

Supplemental Table 1. *C. elegans* MudPIT

Supplemental Table 2. HEK293T MudPIT

## Acknowledgements

We thank the labs of Maurine Linder, Carrie Adler, and Natasza Kurpios for sharing space and reagents. For technical advice and assistance, we thank Wendy Greentree (tissue culture), Jamie Moseley (biochemistry), Ed Partlow (CRISPR reagents and strains), and Maria Henriquez (media prep). We thank Ho Yi Mak and Alejandro Sánchez Alvarado for supporting the identification of the *mlt-4(E407K)* allele, Joe Guinness and Erika Mudrak at the Cornell Statistical Consulting Unit for statistical advice, David Fay, Barth Grant, and Karen Oogema for sharing strains and reagents, Michael Goldberg, Eric Alani, and Carrie Adler for providing constructive criticism that greatly improved the manuscript. UCSF ChimeraX was developed by the Resource for Biocomputing, Visualization, and Informatics at the University of California, San Francisco, with support from National Institutes of Health R01-GM129325 and the Office of Cyber Infrastructure and Computational Biology, National Institute of Allergy and Infectious Diseases. Imaging data was acquired through the Cornell Institute of Biotechnology’s Imaging Facility, with NYSTEM (C029155) and NIH (S10OD018516) funding for the shared Zeiss LSM 880 confocal/multiphoton microscope. Gwendolyn M. Beacham was supported by an NSF graduate research fellowship DGE-1650441. This work was supported by a grant from National Institutes of Health (R01 GM127548-02) awarded to Gunther Hollopeter.

---

**Supplemental Figure 1.**
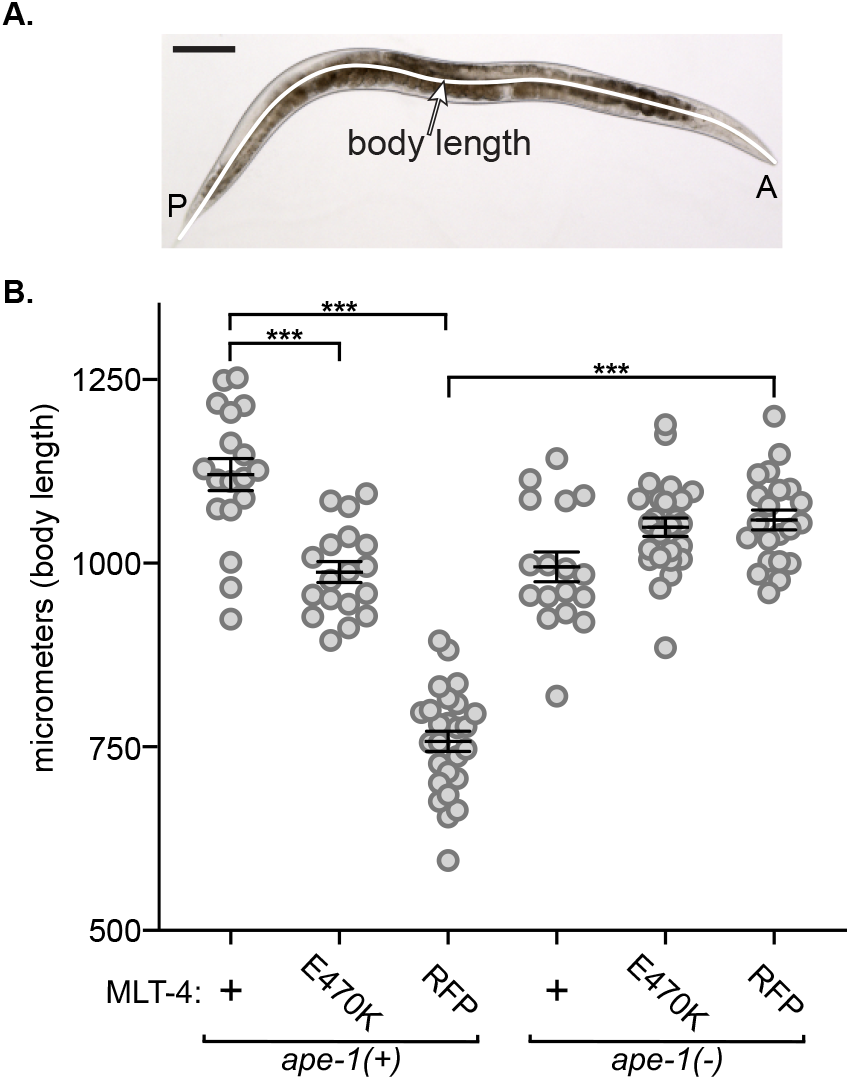
Loss of *ape-1* suppresses the *mlt-4:RFP* dumpy phenotype. **A**. Brightfield image of a wildtype adult worm with an example line drawn for measuring body length. P, posterior; A, anterior. Scale bar is 100 μm. **B**. Data represent body lengths of adult worms, n = 17–26 worms. Bars represent mean ± SEM. ***p<0.0001, ANOVA analysis with Tukey’s post-hoc test. +, wildtype; -, knockout; E470K, single amino acid change; RFP, C-terminal TagRFP:HA-tag.

**Supplemental Figure 2.**
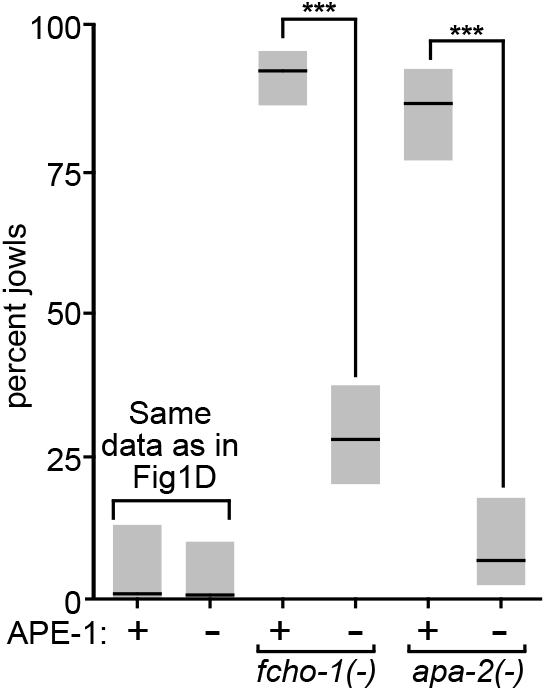
Loss of *ape-1* suppresses jowls of mutants with inactive AP2. Jowls assay. Bars indicate percentage of worms exhibiting jowls, n = 51–126 worms. Gray boxes represent 95% confidence intervals. These data were collected in the same set of strains shown in Figure 1D. The first two samples are repeated from Figure 1D. ***p<0.0001, ANOVA analysis with Tukey’s post-hoc test. +, wildtype; -, knockout.

**Supplemental Figure 3.**
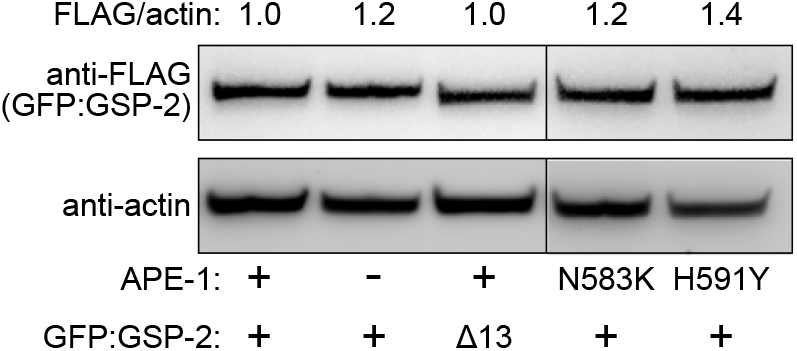
Western blot analysis to detect 3xFLAG epitope on GFP:GSP-2 in whole-worm lysates. Band intensity ratios (above) were normalized to the APE-1(+) GFP:GSP-2(+) ratio (left lane). +, wildtype version; -, knockout; N583K and H591Y, single amino acid changes; Δ13, deletion of the final 13 amino acids. (Note, a MLT-4:HT allele was also present in these strains).

**Supplemental Figure 4.**
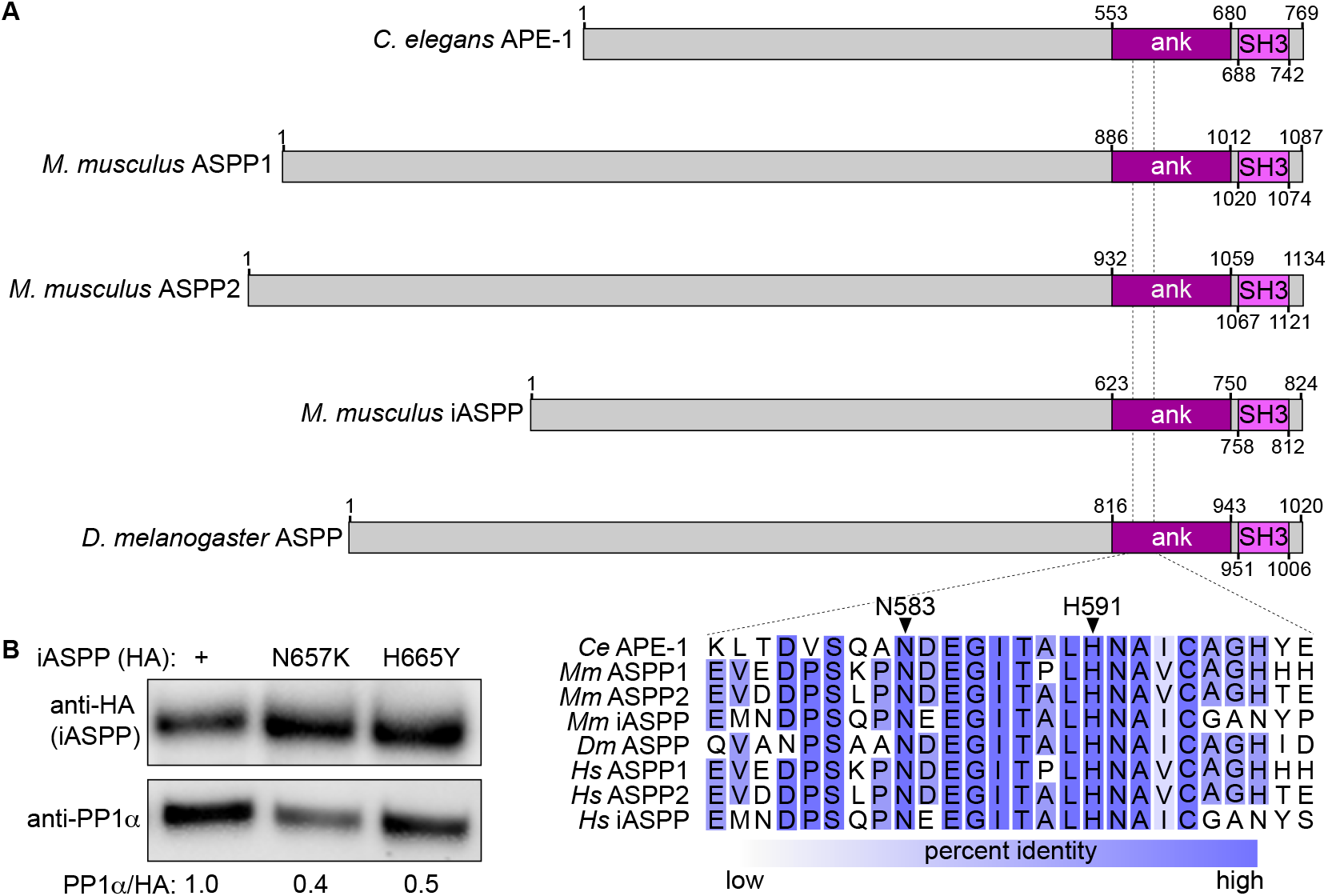
Conservation of the ankyrin repeat missense mutations. **A**. Protein models of APE-1 homologs showing conserved domains. Amino acid numbers are indicated. Dotted lines indicate the residues surrounding worm N583 and H591, aligned below and colored by percent identity. **B**. Pulldown assays using affinity tagged *H. sapiens* iASPP C-terminal domains (amino acids 602-828) from HEK293T cell lysates. iASPP baits had an HA-tag and were wildtype (+) or mutated (N657K and H665Y) as indicated above. Proteins were cleaved from the affinity tag (HaloTag), run on an SDS-PAGE gel, and blotted with anti-HA (top) and anti-PP1*α* (bottom). Intensity ratios of anti-PP1*α* to anti-HA were calculated and normalized to iASPP (+) ratio (bottom values). N657K and H665Y are mutations of the amino acids homologous to worm N583 and H591. ank, ankyrin repeats; SH3, Src Homology 3 domain; *Ce, C. elegans*; *Mm, M. musculus*; *Dm, D. melanogaster; Hs, H. sapiens*.

## References

Alessi, D. et al. (1992) ‘The control of protein phosphatase-1 by targetting subunits. The major myosin phosphatase in avian smooth muscle is a novel form of protein phosphatase-1’, European Journal of Biochemistry, pp. 1023–1035. doi:10.1111/j.1432-1033.1992.tb17508.x.

Beacham, G.M. et al. (2018) ‘NECAPs are negative regulators of the AP2 clathrin adaptor complex’, eLife, 7. doi:10.7554/eLife.32242.

Bergamaschi, D. et al. (2003) ‘iASPP oncoprotein is a key inhibitor of p53 conserved from worm to human’, Nature genetics, 33(2), pp. 162–167.

Bertolotti, A. (2018) ‘The split protein phosphatase system’, Biochemical Journal, 475(23), pp. 3707–3723.

Bertran, M.T. et al. (2019) ‘ASPP proteins discriminate between PP1 catalytic subunits through their SH3 domain and the PP1 C-tail’, Nature communications, 10(1), p. 771.

Bollen, M. et al. (2010) ‘The extended PP1 toolkit: designed to create specificity’, Trends in biochemical sciences, 35(8), pp. 450–458.

Chen, M.J., Dixon, J.E. and Manning, G. (2017) ‘Genomics and evolution of protein phosphatases’, Science signaling, 10(474). doi:10.1126/scisignal.aag1796.

Chisholm, A.D. and Hsiao, T.I. (2012) ‘The Caenorhabditis elegans epidermis as a model skin. I: development, patterning, and growth’, Wiley interdisciplinary reviews. Developmental biology, 1(6), pp. 861–878.

Cohen, P.T.W. (2002) ‘Protein phosphatase 1--targeted in many directions’, Journal of cell science, 115(Pt 2), pp. 241–256.

Cong, W. et al. (2010) ‘ASPP2 regulates epithelial cell polarity through the PAR complex’, Current biology: CB, 20(15), pp. 1408–1414.

Costa, M., Draper, B.W. and Priess, J.R. (1997) ‘The Role of Actin Filaments in Patterning theCaenorhabditis elegansCuticle’, Developmental biology, 184(2), pp. 373–384.

Dedeić, Z. et al. (2018) ‘Cell autonomous role of iASPP deficiency in causing cardiocutaneous disorders’, Cell death and differentiation, 25(7), pp. 1289–1303.

Encell, L.P. (2013) ‘Development of a Dehalogenase-Based Protein Fusion Tag Capable of Rapid, Selective and Covalent Attachment to Customizable Ligands’, Current Chemical Genomics, pp. 55–71. doi:10.2174/1875397301206010055.

Falik-Zaccai, T.C. et al. (2017) ‘Sequence variation in PPP1R13L results in a novel form of cardio-cutaneous syndrome’, EMBO molecular medicine, 9(9), p. 1326.

Florens, L. and Washburn, M.P. (2006) ‘Proteomic analysis by multidimensional protein identification technology’, Methods in molecular biology, 328, pp. 159–175.

Ghanta, K.S. and Mello, C.C. (2020) ‘Melting dsDNA Donor Molecules Greatly Improves Precision Genome Editing in Caenorhabditis elegans’, Genetics, 216(3), pp. 643–650.

Gibson, D.G. et al. (2009) ‘Enzymatic assembly of DNA molecules up to several hundred kilobases’, Nature methods, 6(5), pp. 343–345.

Goddard, T.D. et al. (2018) ‘UCSF ChimeraX: Meeting modern challenges in visualization and analysis’, Protein science: a publication of the Protein Society, 27(1), pp. 14–25.

Grimm, J.B. et al. (2015) ‘A general method to improve fluorophores for live-cell and single-molecule microscopy’, Nature methods, 12(3), pp. 244–50, 3 p following 250.

Gu, M. et al. (2013) ‘AP2 hemicomplexes contribute independently to synaptic vesicle endocytosis’, eLife, 2, p. e00190.

Hadwiger, G. et al. (2010) ‘A monoclonal antibody toolkit for C. elegans’, PloS one, 5(4), p. e10161.

Hattersley, N. et al. (2016) ‘A Nucleoporin Docks Protein Phosphatase 1 to Direct Meiotic Chromosome Segregation and Nuclear Assembly’, Developmental cell, 38(5), pp. 463–477.

Helps, N.R. et al. (1995) ‘Protein phosphatase 1 interacts with p53BP2, a protein which binds to the tumour suppressor p53’, FEBS letters, 377(3), pp. 295–300.

Hendrickx, A. et al. (2009) ‘Docking motif-guided mapping of the interactome of protein phosphatase-1’, Chemistry & biology, 16(4), pp. 365–371.

Herron, B.J. et al. (2005) ‘A mutation in NFkB interacting protein 1 results in cardiomyopathy and abnormal skin development in wa3 mice’, Human molecular genetics, 14(5), pp. 667–677.

Hirashima, M. et al. (2008) ‘Lymphatic vessel assembly is impaired in Aspp1-deficient mouse embryos’, Developmental biology, 316(1), pp. 149–159.

Hollopeter, G. et al. (2014) ‘The membrane-associated proteins FCHo and SGIP are allosteric activators of the AP2 clathrin adaptor complex’, eLife, 3. doi:10.7554/eLife.03648.

Hubbard, M.J. and Cohen, P. (1989) ‘Regulation of protein phosphatase-1G from rabbit skeletal muscle. 2. Catalytic subunit translocation is a mechanism for reversible inhibition of activity toward glycogen-bound substrates’, European journal of biochemistry / FEBS, 186(3), pp. 711–716.

Hubbard, M.J. and Cohen, P. (1993) ‘On target with a new mechanism for the regulation of protein phosphorylation’, Trends in biochemical sciences, 18(5), pp. 172–177.

Iwabuchi, K. et al. (1994) ‘Two cellular proteins that bind to wild-type but not mutant p53’, Proceedings of the National Academy of Sciences of the United States of America, 91(13), pp. 6098–6102.

Joseph, B.B. et al. (2020) ‘Control of clathrin-mediated endocytosis by NIMA family kinases’, PLoS genetics, 16(2), p. e1008633.

Joseph, B.B., Blouin, N.A. and Fay, D.S. (2018) ‘Use of a sibling subtraction method for identifying causal mutations in Caenorhabditis elegans by Whole-genome sequencing’, G3 (Bethesda, Md.), 8(2), pp. 669–678.

Kim, E. et al. (2013) ‘Long-term imaging of Caenorhabditis elegans using nanoparticle-mediated immobilization’, PloS one, 8(1), p. e53419.

Langton, P.F. et al. (2009) ‘The dASPP-dRASSF8 complex regulates cell-cell adhesion during Drosophila retinal morphogenesis’, Current biology: CB, 19(23), pp. 1969–1978.

Lažetić, V. et al. (2018) ‘Actin organization and endocytic trafficking are controlled by a network linking NIMA-related kinases to the CDC-42-SID-3/ACK1 pathway’, PLoS genetics, 14(4), p. e1007313.

Lažetić, V. and Fay, D.S. (2017) ‘Conserved Ankyrin Repeat Proteins and Their NIMA Kinase Partners Regulate Extracellular Matrix Remodeling and Intracellular Trafficking in Caenorhabditis elegans’, Genetics, 205(1), pp. 273–293.

Lienkamp, S., Ganner, A. and Walz, G. (2012) ‘Inversin, Wnt signaling and primary cilia’, Differentiation; research in biological diversity, 83(2), pp. S49–55.

Liu, C.-Y. et al. (2011) ‘PP1 cooperates with ASPP2 to dephosphorylate and activate TAZ’, The Journal of biological chemistry, 286(7), pp. 5558–5566.

Llanos, S. et al. (2011) ‘Inhibitory member of the apoptosis-stimulating proteins of the p53 family (iASPP) interacts with protein phosphatase 1 via a noncanonical binding motif’, The Journal of biological chemistry, 286(50), pp. 43039–43044.

Los, G.V. et al. (2008) ‘HaloTag: a novel protein labeling technology for cell imaging and protein analysis’, ACS chemical biology, 3(6), pp. 373–382.

Mangon, A. et al. (2021) ‘iASPP contributes to cell cortex rigidity, mitotic cell rounding, and spindle positioning’, The Journal of cell biology, 220(12). doi:10.1083/jcb.202012002.

Manning, G. et al. (2002) ‘The protein kinase complement of the human genome’, Science, 298(5600), pp. 1912–1934.

Mello, C.C. et al. (1991) ‘Efficient gene transfer in C.elegans: extrachromosomal maintenance and integration of transforming sequences’, The EMBO journal, 10(12), pp. 3959–3970.

Notari, M. et al. (2015) ‘iASPP, a previously unidentified regulator of desmosomes, prevents arrhythmogenic right ventricular cardiomyopathy (ARVC)-induced sudden death’, Proceedings of the National Academy of Sciences of the United States of America, 112(9), pp. E973–81.

Olsen, J.V. et al. (2006) ‘Global, in vivo, and site-specific phosphorylation dynamics in signaling networks’, Cell, 127(3), pp. 635–648.

Patel, S. et al. (2008) ‘Molecular interactions of ASPP1 and ASPP2 with the p53 protein family and the apoptotic promoters PUMA and Bax’, Nucleic acids research, 36(16), pp. 5139–5151.

Pettersen, E.F. et al. (2021) ‘UCSF ChimeraX: Structure visualization for researchers, educators, and developers’, Protein science: a publication of the Protein Society, 30(1), pp. 70–82.

Ramulu, P. and Nathans, J. (2001) ‘Cellular and Subcellular Localization, N-terminal Acylation, and Calcium Binding of Caenorhabditis elegans Protein Phosphatase with EF-hands’, Journal of Biological Chemistry, pp. 25127–25135. doi:10.1074/jbc.m011712200.

Royer, C. et al. (2014) ‘ASPP2 links the apical lateral polarity complex to the regulation of YAP activity in epithelial cells’, PloS one, 9(10), p. e111384.

Samuels-Lev, Y. et al. (2001) ‘ASPP proteins specifically stimulate the apoptotic function of p53’, Molecular cell, 8(4), pp. 781–794.

Sassa, T. et al. (2003) ‘Role of Caenorhabditis elegans protein phosphatase type 1, CeGLC-7β,in metaphase to anaphase transition during embryonic development’, Experimental cell research, 287(2), pp. 350–360.

Schindelin, J. et al. (2012) ‘Fiji: an open-source platform for biological-image analysis’, Nature methods, 9(7), pp. 676–682.

Sharma, K. et al. (2014) ‘Ultradeep human phosphoproteome reveals a distinct regulatory nature of Tyr and Ser/Thr-based signaling’, Cell reports, 8(5), pp. 1583–1594.

Shi, Y. (2009) ‘Serine/threonine phosphatases: mechanism through structure’, Cell, 139(3), pp. 468–484.

Simpson, M.A. et al. (2009) ‘A mutation in NFkappaB interacting protein 1 causes cardiomyopathy and woolly haircoat syndrome of Poll Hereford cattle’, Animal genetics, 40(1), pp. 42–46.

Skene-Arnold, T.D. et al. (2013) ‘Molecular mechanisms underlying the interaction of protein phosphatase-1c with ASPP proteins’, Biochemical Journal, 449(3), pp. 649–659.

Sottocornola, R. et al. (2010) ‘ASPP2 binds Par-3 and controls the polarity and proliferation of neural progenitors during CNS development’, Developmental cell, 19(1), pp. 126–137.

Sullivan, A. and Lu, X. (2007) ‘ASPP: a new family of oncogenes and tumour suppressor genes’, British journal of cancer, 96(2), pp. 196–200.

Tabb, D.L., McDonald, W.H. and Yates, J.R., 3rd (2002) ‘DTASelect and Contrast: tools for assembling and comparing protein identifications from shotgun proteomics’, Journal of proteome research, 1(1), pp. 21–26.

Tanaka, J. et al. (1998) ‘Interaction of myosin phosphatase target subunit 1 with the catalytic subunit of type 1 protein phosphatase’, Biochemistry, 37(47), pp. 16697–16703.

Terrak, M. et al. (2004) ‘Structural basis of protein phosphatase 1 regulation’, Nature, 429(6993), pp. 780–784.

Tidow, H. et al. (2006) ‘Effects of oncogenic mutations and DNA response elements on the binding of p53 to p53-binding protein 2 (53BP2)’, The Journal of biological chemistry, 281(43), pp. 32526–32533.

Toonen, J., Liang, L. and Sidjanin, D.J. (2012) ‘Waved with open eyelids 2 (woe2) is a novel spontaneous mouse mutation in the protein phosphatase 1, regulatory (inhibitor) subunit 13 like (Ppp1r13l)gene’, BMC Genetics. doi:10.1186/1471-2156-13-76.

Traweger, A. et al. (2008) ‘Protein phosphatase 1 regulates the phosphorylation state of the polarity scaffold Par-3’, Proceedings of the National Academy of Sciences of the United States of America, 105(30), pp. 10402–10407.

tzw-wen (2022) tzw-wen/kite: doi:10.5281/zenodo.5914885.

Varland, S., Vandekerckhove, J. and Drazic, A. (2019) ‘Actin Post-translational Modifications: The Cinderella of Cytoskeletal Control’, Trends in biochemical sciences, 44(6), pp. 502–516.

Vives, V. et al. (2006) ‘ASPP2 is a haploinsufficient tumor suppressor that cooperates with p53 to suppress tumor growth’, Genes & development, 20(10), pp. 1262–1267.

Washburn, M.P., Wolters, D. and Yates, J.R., 3rd (2001) ‘Large-scale analysis of the yeast proteome by multidimensional protein identification technology’, Nature biotechnology, 19(3), pp. 242–247.

Waterhouse, A.M. et al. (2009) ‘Jalview Version 2—a multiple sequence alignment editor and analysis workbench’, Bioinformatics, 25(9), pp. 1189–1191.

Whittington, R.J. and Cook, R.W. (1988) ‘Cardiomyopathy and woolly haircoat syndrome of Poll Hereford cattle: electrocardiographic findings in affected and unaffected calves’, Australian veterinary journal, 65(11), pp. 341–344.

Wloga, D., Joachimiak, E. and Fabczak, H. (2017) ‘Tubulin Post-Translational Modifications and Microtubule Dynamics’, International journal of molecular sciences, 18(10). doi:10.3390/ijms18102207.

Wu, J.-C. et al. (2012) ‘Sperm development and motility are regulated by PP1 phosphatases in Caenorhabditis elegans’, Genetics, 190(1), pp. 143–157.

Xu, T. et al. (2015) ‘ProLuCID: An improved SEQUEST-like algorithm with enhanced sensitivity and specificity’, Journal of proteomics, 129, pp. 16–24.

Zhang, Y. et al. (2010) ‘Refinements to label free proteome quantitation: how to deal with peptides shared by multiple proteins’, Analytical chemistry, 82(6), pp. 2272–2281.

Zhang, Y. et al. (2011) ‘Improving proteomics mass accuracy by dynamic offline lock mass’, Analytical chemistry, 83(24), pp. 9344–9351.

Zhou, Y. et al. (2019) ‘Flexible Tethering of ASPP Proteins Facilitates PP-1c Catalysis’, Structure, 27(10), pp. 1485–1496.e4.

